# Late-Life Alcohol Exposure Does Not Exacerbate Age-Dependent Reductions in Mouse Spatial Memory and Brain TFEB Activity

**DOI:** 10.1101/2024.02.23.581774

**Authors:** Hao Chen, Kaitlyn Hinz, Chen Zhang, Yssa Rodriguez, Sha Neisha Williams, Mengwei Niu, Xiaowen Ma, Xiaojuan Chao, Alexandria L. Frazier, Kenneth E. McCarson, Xiaowan Wang, Zheyun Peng, Wanqing Liu, Hong-Min Ni, Jianhua Zhang, Russell H. Swerdlow, Wen-Xing Ding

**Author notes:** Correspondence to: Wen-Xing Ding, Department of Pharmacology, Toxicology and Therapeutics; The University of Kansas Medical Center; MS 1018; 3901 Rainbow Blvd. Kansas City, KS 66160.

## Abstract

Alcohol consumption is believed to affect Alzheimer’s disease (AD) risk, but the contributing mechanisms are not well understood. A potential mediator of the proposed alcohol-AD connection is autophagy, a degradation pathway that maintains organelle and protein homeostasis. Autophagy is in turn regulated through the activity of Transcription factor EB (TFEB), which promotes lysosome and autophagy-related gene expression. To explore the effect of alcohol on brain TFEB and autophagy, we exposed young (3-month old) and aged (23-month old) mice to two alcohol-feeding paradigms and assessed biochemical, transcriptome, histology, and behavioral endpoints. In young mice, alcohol decreased hippocampal nuclear TFEB staining but increased SQSTM1/p62, LC3-II, ubiquitinated proteins, and phosphorylated Tau. Hippocampal TFEB activity was lower in aged mice than it was in young mice, and Gao-binge alcohol feeding did not worsen the age-related reduction in TFEB activity. To better assess the impact of chronic alcohol exposure, we fed young and aged mice alcohol for four weeks before completing Morris Water and Barnes Maze spatial memory testing. The aged mice showed worse spatial memory on both tests. While alcohol feeding slightly impaired spatial memory in the young mice, it had little effect or even slightly improved spatial memory in the aged mice. These findings suggest that aging is a far more important driver of spatial memory impairment and reduced autophagy flux than alcohol consumption.

## Introduction

Alzheimer’s disease (AD) is a prevalent form of age-dependent progressive dementia that affects more than 4% of individuals aged 65 years and above. The primary pathological characteristics of AD are the buildup of extracellular β-amyloid plaques and intracellular fibrillary tangles, which are composed of aggregated hyperphosphorylated tau (Hardy 2006). In addition to the accumulation of protein aggregates, mitochondrial dysfunction also plays a critical role in the pathogenesis of AD and perhaps some AD related disorders (ADRD) (Swerdlow 2018, Van Giau, An et al. 2018). Extensive clinical studies strongly support a link between alcohol use disorders with AD and ADRD (Stavro, Pelletier et al. 2013, Piumatti, Moore et al. 2018). Heavy alcohol drinkers have impaired cognitive function even in the absence of a dementia diagnosis. Individuals with alcohol use disorders also have a 2-fold greater risk of developing AD versus the general population (Schwarzinger, Pollock et al. 2018). However, some epidemiology studies also suggest that moderate drinking is beneficial and decrease the risk of dementia (Chan, Chiu et al. 2010, Jeon, Han et al. 2023). These divergent studies may reflect inconsistencies in alcohol and AD measurement parameters. However, the molecular mechanisms underlying the effects of alcohol on brain function and the pathogenesis of AD remain largely unknown.

Autophagy is a highly conserved lysosomal degradation pathway that is activated in response to stress, such as deprivation of nutrients or growth factors, as a survival mechanism (Klionsky and Emr 2000, Mizushima, Levine et al. 2008). Autophagy can help remove excess protein aggregates (aggrephagy) or damaged organelles such as mitochondria (mitophagy) to protect against cell death (Singh, Kaushik et al. 2009, Czaja, Ding et al. 2013, Williams, Ni et al. 2015, Martini-Stoica, Xu et al. 2016). The lysosome is the terminal component of autophagy and contains more than 50 acid hydrolases. The transcriptional regulation of genes that facilitate lysosome biogenesis and autophagy play a critical role in autophagy (Settembre, Fraldi et al. 2013). Transcription factor EB (TFEB) is a basic helix-loop-helix leucine zipper transcription factor belonging to the coordinated lysosomal expression and regulation (CLEAR) gene network (Settembre, Fraldi et al. 2013). TFEB is a master regulator for transcription of lysosome biogenesis and autophagy genes (Settembre, Di Malta et al. 2011, Settembre, Zoncu et al. 2012). In response to an increased need for autophagic degradation, TFEB coordinates an efficient transcriptional program that upregulates the expression of genes that are responsible for both the early (autophagosome formation) and late (lysosome biogenesis) phases of autophagy. SIRT1, an NAD+-dependent deacetylase, deacetylates TFEB at lysine 116 and this induces a TFEB-mediated lysosomal biogenesis that promotes the degradation of fibrillar Aβ by microglia (Bao, Zheng et al. 2016).

We and others have demonstrated that alcohol consumption impairs TFEB-mediated lysosomal biogenesis in liver and pancreas, which promotes alcoholic hepatitis and pancreatitis (Chao, Wang et al. 2018, Babuta, Furi et al. 2019, Wang, Ni et al. 2020). Whether and how alcohol affects TFEB in the brain, and especially in the hippocampi that are affected early in AD, has not been studied.

In addition to alcohol abuse, increased age is a well-known risk factor for AD. Both autophagy and lysosomal functions decline with aging (Levine and Kroemer 2019). It has been demonstrated that brains from individuals with AD have impaired autophagy. This results in excessive accumulation of immature autophagosomes, which is caused by defects in the fusion of autophagosomes with lysosomes or hindered retrograde traffic of autophagosomes towards the neuronal cell body (Nixon 2007). In AD, a buildup of immature autophagosomes can cause inadequate autophagy, leading to the accumulation of tau and β-amyloid plaques. A decrease in nuclear TFEB levels has been observed in the subcellular fractionation analysis of AD patient brains. There is a strong inverse correlation between hippocampal nuclear TFEB levels and the severity of AD pathology (Wang, Wang et al. 2016). Moreover, decreased TFEB function has been observed in AD patient lymphocytes and monocytes, which may migrate to damaged central nervous system (CNS) regions and regulate AD progression (Tiribuzi, Crispoltoni et al. 2014). Furthermore, expression levels of the CLEAR gene network also decreased in cultured fibroblasts from AD patients and iPSC-derived human AD neurons (Reddy, Cusack et al. 2016). More importantly, overexpression of TFEB promotes the clearance of aberrant tau proteins in the rTg4510 and P301S mouse models of tauopathy (Polito, Li et al. 2014, Wang, Wang et al. 2016). More recently, the curcumin analog C1 was shown to activate TFEB and increases lysosome biogenesis, which in turn decreased beta-amyloid precursor protein and Tau pathology in three AD animal models (5XFAD, P301S and 3XTg-AD) (Song, Malampati et al. 2020).

In the present study we used chronic plus and binge alcohol models to explore how alcohol affects autophagy-lysosome related pathways in mouse brains. We also used a chronic alcohol feeding model in young and aged mice to investigate whether age and alcohol affected mouse cognitive performance. Our results show that aging and alcohol impair TFEB signaling in the brains of young mice, and in old mice aging more robustly affects cognitive performance than alcohol.

## Materials and Methods

### Animal Experiments

Male 3- and 23-month-old C57BL/6N mice were obtained from Charles River Laboratories (Wilmington, MA) and housed at the AAALAC-accredited facility at KUMC for at least one week on regular chow diet for acclimation. All mice were specific pathogen free (SPF) and received human care in a barrier rodent facility under standard experimental conditions. Animal studies were approved and performed under the of Institutional Animal Care and Use Committee (IACUC) at the University of Kansas Medical Center (Kansas City, KS).

Subsequently, for the Gao-binge alcohol model, mice were acclimated to the Lieber-DeCarli liquid control diet (F1259SP, Bio-Serv) for 5 days followed by further feeding with the liquid control or ethanol diet (F1258SP, Bio-Serv, 5% ethanol) for 10 days. The volume of control diet given to mice was matched to the volume of ethanol diet consumed. On the last day of feeding, mice were further given 5 g/kg ethanol or 9 g/kg maltose dextran and sacrificed 8 hours later. On the last day of feeding, mice were euthanized 8 hours after the gavage. Brain tissues and blood samples were collected. Left-hemi brains were collected for western blotting and right-hemi brains were obtained for cryosection, hematoxylin and eosin (H&E) staining as well as confocal microscopy.

Prior to behavioral testing, mice were acclimated to the Lieber-DeCarli liquid control diet) for 5 days followed by further feeding with the liquid control or ethanol diet (5% ethanol) for 4 weeks. After the last day of alcohol and liquid control diet feeding, mice were returned to chow diet for recovery of 3 days followed by either Morris Water Maze or Barnes Maze testing.

### Mouse behavioral testing

The Morris Water Maze test was used to assess spatial learning and memory using well-defined protocols (Morris, Garrud et al. 1982, Vorhees and Williams 2006). Mice were allowed to acclimate to the testing environment for at least one hour before testing. Mice were randomized by diet treatment group, to which the experimenter was blinded during testing. In brief, mice learn to find a hidden platform (20 cm diameter) in an open swimming pool (200 cm diameter) filled with 21 °C water, which was made opaque with nontoxic poster paint. A digital camera suspended above the maze recorded the animal’s position during the test. Four trials were performed each day for five days. Each trial started at a different position (N, E, SE, NW) while the platform was kept in a fixed location (NE). Each trial lasted 60 seconds, followed by 30 seconds during which mice were allowed to remain on the platform to strengthen their memory of platform location. After five days of memory acquisition trials, mice were subjected to a probe (retention) trial in which the platform was removed. The latency to reaching the platform, the time they spent in the target quadrant, and the number of times they passed the previous platform location were analyzed by using the animal tracker plugin of ImageJ (National Institutes of Health).

A Barnes Maze testing protocol was adapted from previous descriptions (Barnes 1979, Pitts 2018, Rodriguez Peris, Scheuber et al. 2024). The memory acquisition protocol consisted of one acclimation session followed by five training sessions. The mice were initially acclimated to the maze by placing them within a goal box located outside of the Barnes maze table for 2 minutes. Goal box acclimation preceded the first training session by at least one day. During the first training session, the mice were placed in the center of the Barnes maze platform covered by an inverted paper container for approximately 20 seconds, and then released to find the goal box. If the mouse failed to find the goal within 3 minutes, it was gently guided to it and kept there for 1 minute before being returned to the home cage. This training was repeated for five consecutive days with four trials occurring each day per mouse. Mice were given intervals of no less than 5 minutes between each trial. The trials were video recorded and analyzed using EthoVision XT (Version 17.0.1630, Noldus Information Technology, Wageningen, The Netherlands).

### RNAseq data analysis

Total RNA was extracted from mouse hippocampi using TRIzol reagent (15596-026; Ambion, ThermoFisher Scientific) and was reverse-transcribed into cDNA using RevertAid Reverse Transcriptase (EP0442; Fermentas, ThermoFisher Scientific). A detailed RNA sequencing analysis was performed as described previously (Ma, Chen et al. 2023). The RNAseq data were analyzed using our previously established technical pipeline. Briefly, we used HISAT2 v.2.1.0.13 to map the high-quality reads to the mouse reference genome (GRCm38.90). Quantification of gene expression was generated with HTSeq-counts v0.6.0. Significant differentially expressed genes (DEGs) were identified with the R package DESeq2. Multiple comparisons were taken into account by converting P values to the Benjamini-Hochberg’s false discovery rate (FDR).

Volcano plots were prepared with R software Version 4.2.0 (https://www.r-project.org/). Pathway analysis was performed using the online tool ShinyGO v0.741. We analyzed the enrichment of up-and down-regulated genes among each EtOH group as compared to the control group. Genes with a log2 fold change >1 or < −1 with an FDR-corrected p value <0.05 were used to perform the analysis. Heatmap analysis based on Gene Ontology (GO) and Kyoto Encyclopedia of Genes and Genomes (KEGG) enrichment analyses and circos plots of the DEGs were analyzed using Metascape (http://metascape.org)

### Western blot analysis

Total brain hippocampi or cortex lysates were prepared in radioimmunoprecipitation assay buffer [1% NP-40, 0.5% sodium deoxycholate, and 0.1% sodium dodecyl (lauryl) sulfate]. Protein (20 µg) was separated by a 12% SDS-PAGE gel, then transferred to a polyvinylidene difluoride (PVDF) membrane. After blocking in TBS buffer (20 mM Tris-HCl, 150 mM sodium chloride) containing 5% (wt/vol) nonfat dry milk for 1 h at room temperature, the membranes were then probed with primary and secondary antibodies, followed by development with Super Signal West Pico chemiluminescent substrate (34080; Life Technologies, Carlsbad, CA). Data analysis was performed by Image lab 6.1 (Bio-Rad, Hercules, CA). The following primary antibodies were used: mouse anti-p62 (Abnova, H00008878-M01, 1:10000), mouse anti-phospho-Tau (ser396) (Cell Signaling, 9632S, 1:3000), rabbit anti-phospho-Tau (ser404) (Cell Signaling, 20194S, 1:2000), mouse anti-total-Tau (Cell Signaling, 4019S, 1:3000), rabbit anti-TFEB (Bethyl, A303-673A, 1:1000), mouse anti-β-actin (Sigma, A5411, 1:10000), mouse anti-PSD95 (Abcam, ab2723, 1:1000), rabbit anti-Synapsin 1 (Millipore, AB1543, 1:1000), mouse anti-ubiquitin(Santa Cruz, sc-8017, 1:3000), and rabbit anti-GAPDH (Cell Signaling, 2118, 1:5000). The following secondary antibodies were used: HRP-goat anti-Mouse IgG (Jackson, 115-035-146, 1:2000), HRP-goat anti-Rabbit IgG (Jackson, 111-035-144, 1:2000).

### Histology and Immunohistochemistry (IHC) Staining

Paraffin-embedded brain sections were stained with hematoxylin and eosin. Additionally, immunostaining with indicated antibodies was performed as previously described. Briefly, paraffin-embedded tissue sections were incubated with primary antibody at 4°C overnight after deparaffinization and heat-induced antigen retrieval in citrate buffer. Sections were then washed and incubated with secondary antibody for 1h at 37°C and developed with ImmPACT Nova RED HRP substrate (Vector Labs, SK-4805). Tissues were counterstained with hematoxylin. The following antibodies were used: rabbit anti-TFEB (Bethyl, A303-673A, 1:1000), and HRP Horse Anti-Rabbit IgG (Vector, MP7401, one drop).

### Electron microscopy analysis

Male mice aged 3 and 23 months were fed with the Gao-binge alcohol model. Their hippocampal tissues were fixed with 2% glutaraldehyde in 0.1 M phosphate buffer (pH 7.4), then treated with 1% OsO4. Thin sections were cut and stained with uranyl acetate and lead citrate. Digital images were taken with a JEM 1016CX electron microscope.

### Immunofluorescence Staining

Brain frozen sections from 3 and 24-months old mice with or without ethanol treatment were used to perform immunostaining. Antigen retrieval was done by boiling the samples in citrated buffer for 15 minutes, followed by blocking with 5% BSA/PBS and 0.5% triton-X for 1 h at room temperature (RT). Slides were mounted in PermaFluor Aqueous Mounting Medium (Thermo Scientific, TA-030-FM) followed by confocal microscopy. The following primary antibodies were used: mouse anti-total-Tau (Cell Signaling, 4019S, 1:3000) and rabbit anti-TFEB (Bethyl, A303-673A, 1:300). The following secondary antibodies were used: Alex488 goat anti-mouse (Jackson, 115-545-146, 1:600) and Alex594 goat anti-rabbit (Jackson, 111-505-144, 1:600).

### Statistical analysis

Statistical comparisons were performed using GraphPad Prism 8 software (https://www.graphpad.com/scientific-software/prism/). No randomization was performed, but data collection and analysis were conducted blindly to the conditions of the experiments by researchers who were blinded to group assignment. Two-way ANOVA followed by Bonferroni post hoc analysis or unpaired Student’s t-test followed by Bonferroni post hoc analysis were applied for data analysis. Numbers of replicates and p values are stated in each figure legend. All data are expressed as means ± SEM. Significance is indicated by the following symbols: \**p* < 0.05, \*\**p* < 0.01, \*\*\**p* < 0.001.

## Results

### Aging and chronic plus binge EtOH (Gao-binge) feeding impair TFEB-mediated autophagy in mouse brains

We first determined the changes of TFEB and several autophagy-related markers in alcohol fed young and aged mouse hippocampi. Results from IHC staining revealed that alcohol feeding decreased nuclear TFEB and increased ubiquitin staining in young mouse hippocampi (**Figure 1 A-B**). Aged mice had decreased nuclear TFEB and increased ubiquitin staining regardless of alcohol feeding (**Figure 1A-B**). Consistent with the IHC results, western blot analysis showed decreased TFEB but increased levels of ubiquitin in alcohol-fed young mouse hippocampi. Aged mice had decreased nuclear TFEB and increased ubiquitin staining, and alcohol feeding did not further alter this finding (**Figure 1C**). Gao-binge alcohol feeding did not appreciably alter p62 but slightly increased LC3-II levels in young mouse hippocampi. The p62 levels were also comparable between young and aged mice regardless of alcohol feeding. However, the levels of LC3-II increased in control diet fed aged mouse hippocampi and further increased with alcohol feeding (**Figure 1C-D**).

**Figure 1.**
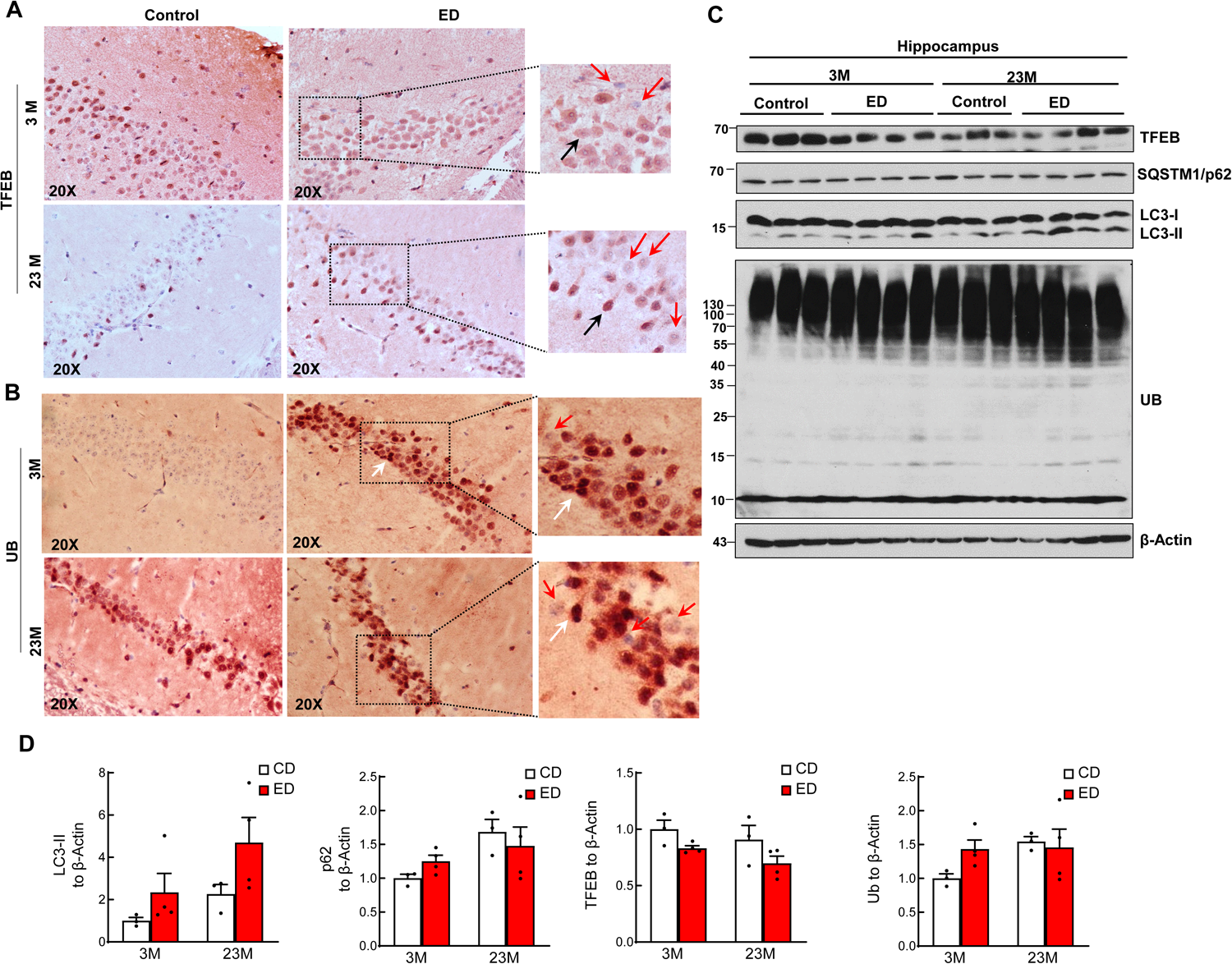
Aging and chronic plus binge EtOH (Ga-binge) feeding impair TFEB-mediated autophagy in mouse brains. Male 3-month-old young mice (3M) and 23-month-old aged mice (23M) were subjected to Gao-binge alcohol feeding. Representative images of immunohistochemistry staining for TFEB **(A)** and ubiquitin (UB) (**B**) are shown. Arrows denotes positive TFEB and UB stained cells. **(C)** Total lysates from hippocampus were subjected to western blot analysis. (**D**) Densitometry analysis from (**C**). Data are presented as means ± SEM (N=3-4). ED: ethanol diet with binge.

Accumulating Aβ fibrils and lipofuscin are often found in late endosomes/lysosomes of AD patient brains (Androuin, Thierry et al. 2022). EM ultrastructure analysis revealed accumulation of abnormal lysosome structures enveloping lipid droplets (LD) and other electron dense materials as well as inclusion body-like structures reminiscent of lipofuscin granules and Aβ fibril-like structures in alcohol fed but not control diet-fed young mouse hippocampi (**Figure 2A-B**). Aged mice had increased lysosomes containing lipofuscin regardless of alcohol feeding (**Figure 2C-D**). Moreover, swollen mitochondria with abnormal cristae were readily detected in aged mouse hippocampi regardless of alcohol feeding (**Figure 2C-D**). Together, these results suggest that aging and chronic plus binge alcohol feeding impair TFEB-mediated autophagy and lead to the accumulation of abnormal lysosome structures containing lipofuscin and Aβ fibril-like structures. Beyond this, the concomitant presence of both aging and alcohol did not demonstrate an additive or synergistic interaction.

**Figure 2.**
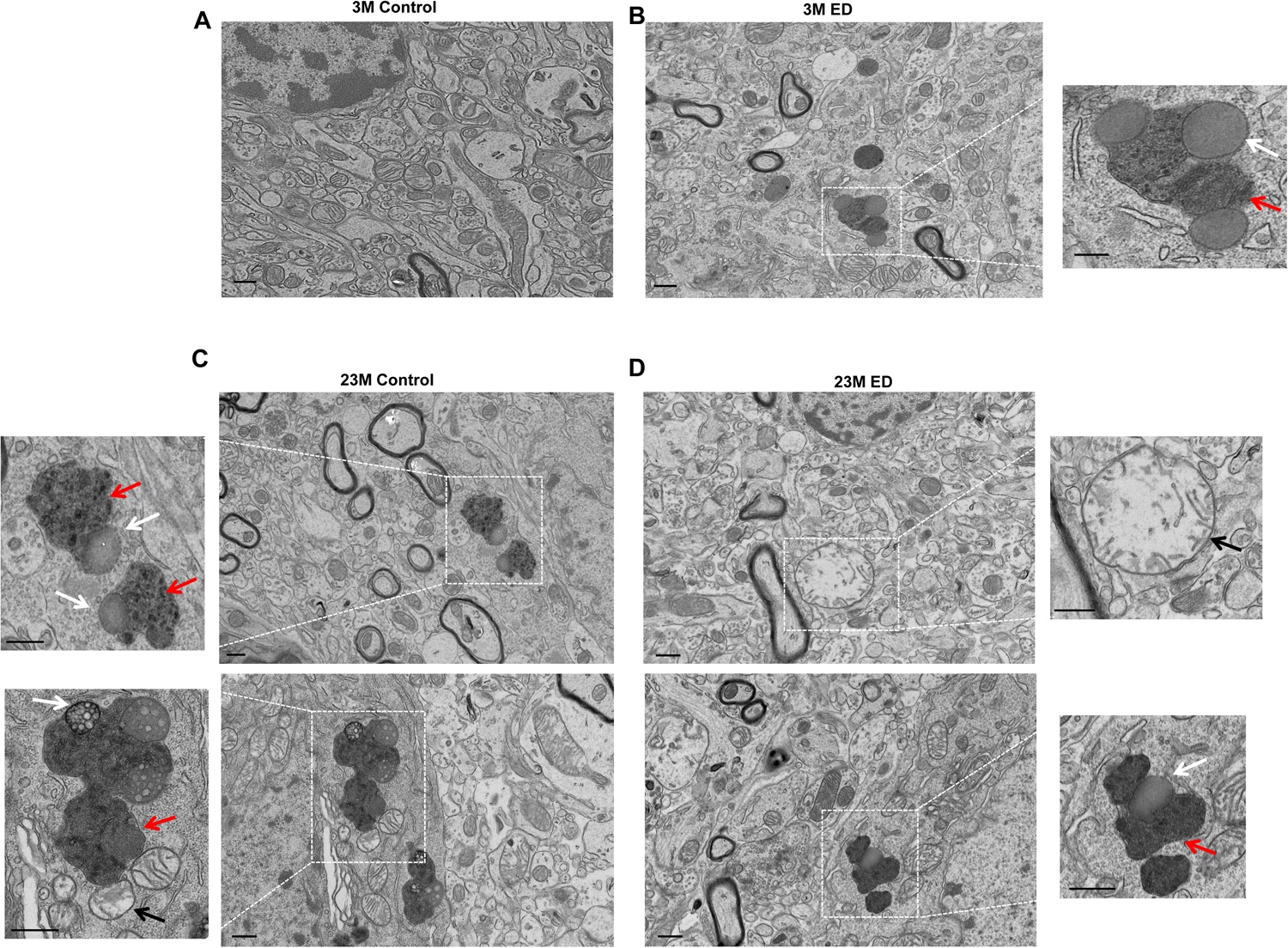
Electron microscopy analysis of ultrastructure of Gao-binge alcohol-fed young and aged mouse hippocampi. Male 3-month-old young mice (3M) and 23-month-old aged mice (23M) were subjected to Gao-binge alcohol feeding. Representative EM images of mouse hippocampi from 3M control (A), ED (B), 23M control (C) and 23M ED (D) mice. Right panels are enlarged images from the boxed area in (B) and (D). Left panels are enlarged images from (C). Red arrows denote lysosomes; white arrows denote lipid droplets in lysosomes, and black arrows denote damaged mitochondria. Bar: 500 nm.

### Transcriptomic analysis of hippocampi from chronic plus binge EtOH (Gao-binge) fed young and aged mice

As shown in the Circos plot, 171 genes were significantly upregulated. In contrast, only 41 genes were significantly downregulated in control diet-fed aged mice compared with control diet-fed young mice (**Figure 3A-B**). Alcohol-fed young mice had 275 genes upregulated, but 244 genes were downregulated compared with control diet-fed young mice. In contrast, alcohol-fed aged mice had 183 upregulated genes, but 284 genes were downregulated compared with control diet-fed aged mice (**Figure 3A-B**). Among the upregulated genes, 169 genes were exclusively upregulated in control diet-fed aged mice compared with control diet-fed young mice, 115 genes in alcohol-fed aged mice compared with control diet-fed aged mice, and 148 genes in alcohol-fed young mice compared with control diet-fed young mice, respectively (**Figure 3A**). Among the commonly upregulated genes, 59 genes were found between “23M CD vs 3M CD” and “3M ED vs 3M CD”, suggesting these genes may be induced either by aging alone or by alcohol in young mice. In contrast, 62 genes were shared between “23M ED vs 23M CD” and “3M ED vs 3M CD”, suggesting these genes were mainly affected by alcohol independent of age (**Figure 3A**).

**Figure 3.**
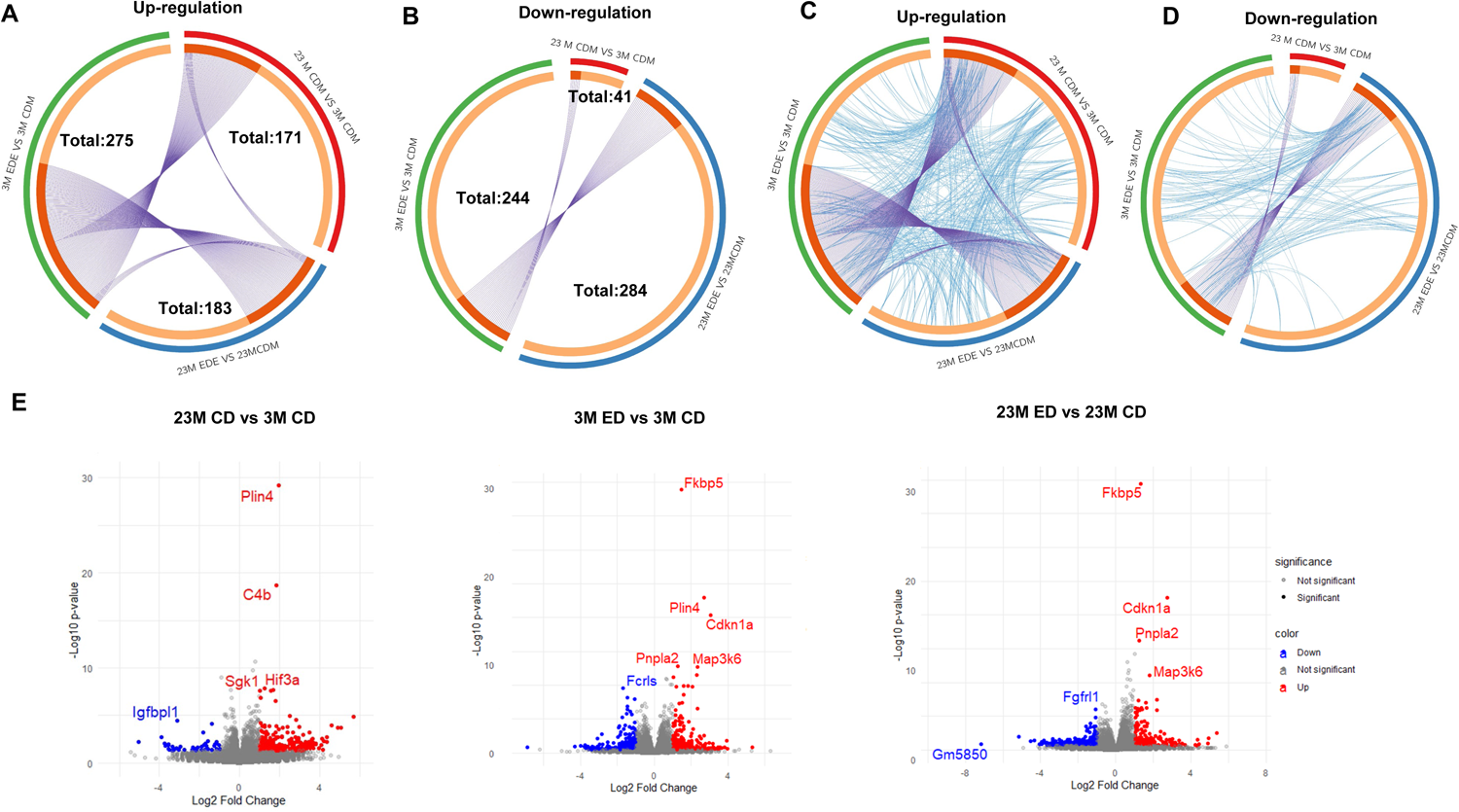
Transcriptomic analysis of Gao-binge alcohol fed young and aged mouse hippocampi. Male 3-month-old young mice (3M) and 23-month-old aged mice (23M) were subjected to Gao-binge alcohol feeding. mRNAs were extracted from the mouse hippocampi after the feeding and subjected to RNAseq analysis. (A-D) Circos plot analysis from the RNAseq data set. Red: 23M CD (control diet) vs 3M CD; blue: 23M ED vs 23M CD, green: 3M ED vs 3M CD. The inner circle represents gene lists, where hits are arranged along the arc. Purple curves link identical genes. Genes that hit multiple lists are colored in dark orange, and genes unique to a list are shown in light orange. Each arc represents a distinct group of genes, color-coded to align with the corresponding experimental conditions, while the interconnecting chords depict shared ontology terms. including the shared term level, where blue curves link genes that belong to the same enriched ontology term. The thickness of each chord is proportional to the number of genes that contribute to the common functional annotation analysis(C) for up-regulated genes (D) for down regulated genes. (E) Volcano plots illustrating differential gene expression across experimental conditions. The x-axis represents the log2 fold change, and the y-axis represents the -log10 p-value, indicating the magnitude and statistical significance of gene expression changes, respectively. Upregulated genes are marked in red, downregulated genes in blue, and non-significant changes in gray. Key genes with a fold change greater than 2 and a p-value less than 0.05 are labeled.

Among the commonly downregulated genes, 33 genes were found exclusively downregulated in control diet-fed aged mice compared with control diet-fed young mice, 250 genes in alcohol-fed aged mice compared with control diet-fed aged mice, and 202 genes in alcohol-fed young mice compared with control diet-fed young mice, respectively (**Figure 3B**). Notably, the intersection comparison of “23M CD vs 3M CD” and “23M ED vs 23M CD”’ yielded eight commonly downregulated genes (*Cenpf, Gm13690, Gm26526, Gm3756, Gm43339, Gm6170, Rpsa-ps10, and Top2a*), indicating a potential overlap in regulatory mechanisms in alcohol and aging although only in a small subset of genes. However, there were 34 common genes downregulated between “23M ED vs 23M CD” and “3M ED vs 3M CD”, suggesting a subset of downregulated genes were affected by alcohol independent of age (**Figure 3B**). Interestingly, there were no shared downregulated genes between the “23M CD vs 3M CD” and “23M ED vs 23M CD” groups (**Figure 3B**), demonstrating no overlap of downregulated genes between alcohol and aging.

In our next comprehensive analysis, we further refined gene set overlaps by incorporating functional annotations, revealing intricate networks of shared biological processes and pathways. **Figure 3C and D** present a chord diagram that illustrates the functional interconnections between the genes either upregulated or downregulated in the three experimental comparison groups (23M CD vs 3M CD, 23M ED vs 23M CD, and 3M ED vs 3M CD). The diagram accentuated the enriched ontology terms, specifically those encompassing fewer than 100 genes to preserve specificity, thus excluding overly general annotations. Each arc represents a distinct group of genes, color-coded to align with the corresponding experimental conditions, while the interconnecting chords depict shared ontology terms. The thickness of each chord is proportional to the number of genes that contribute to the common functional annotation, highlighting the most significantly enriched pathways. The biological systems under study exhibit a complex network of gene regulation, with certain pathways being distinctly modulated (either upregulated or downregulated) across different conditions. Compared to the down-regulated gene network, the up-regulated gene network appears to be more interconnected, suggesting a more complex and potentially more coordinated response. This may imply that the up-regulated genes are part of a robust activation process, possibly related to a stress response or other significant cellular changes in the combination of alcohol feeding and aging.

Volcano plot analysis revealed distinct gene expression profiles among the experimental conditions (**Figure 3E**). Notably, genes such as *Plin4*, which is responsible for lipid droplet regulation, *Hif3a*, a transcription factor that regulates gene expression in response to low oxygen, and *Fmo2*, a flavin-containing monooxygenase, showed significant upregulation in the 23M CD group compared to the 3M CD group. These findings suggest that aging may increase lipid and xenobiotic metabolism and cause hypoxia in mouse brains. It was observed that both *Plin4* and *Pnpla2* genes showed higher expression in the 3M ED group compared to the 3m CD group. However, among the 23M ED and 23M CD groups, only *Pnpla2* was found to have higher expression, while *Plin4* did not show a significant difference. This suggests that alcohol may cause general lipid metabolism changes regardless of age. Notably, *Fkbp5* (FK506-binding protein 5) was identified as the highest expression gene in both the 23M ED and 3M ED groups but not in the 23M CD group. This suggests that *Fkbp5* may be specifically induced by alcohol. FKBP5 is a key molecule in the stress response and a particular single nucleotide polymorphism (SNP) has been implicated in the hypothalamic-pituitary-adrenal (HPA) axis and the development of stress-related psychiatric disorders such as posttraumatic stress disorder (PTSD), as well as the pathophysiology of stress-related disorders and AD (Fujii, Hori et al. 2014, Fujii, Ota et al. 2014).

A heatmap analysis of the RNAseq data showed that alcohol metabolism genes such as *Ahd, Aldh*, and *Cyp2e1* increased in alcohol-fed mice regardless of age (**Figure 4A**). On the other hand, the expression of inflammatory genes was generally higher in aged mice compared to young mice, regardless of alcohol consumption (**Figure 4B**). Additionally, some of the TFEB target genes including *Atpv1a, Atpv0c, Atpv0b, Atpv1h* and autophagy-related genes (*Atg4b, Atg12, Wipi2, Atg4c, Atg4d, Pik3c3, Vps18*) were downregulated in aged mice as compared to young mice, with alcohol consumption having little effect (**Figure 4 C-D**).

**Figure 4.**
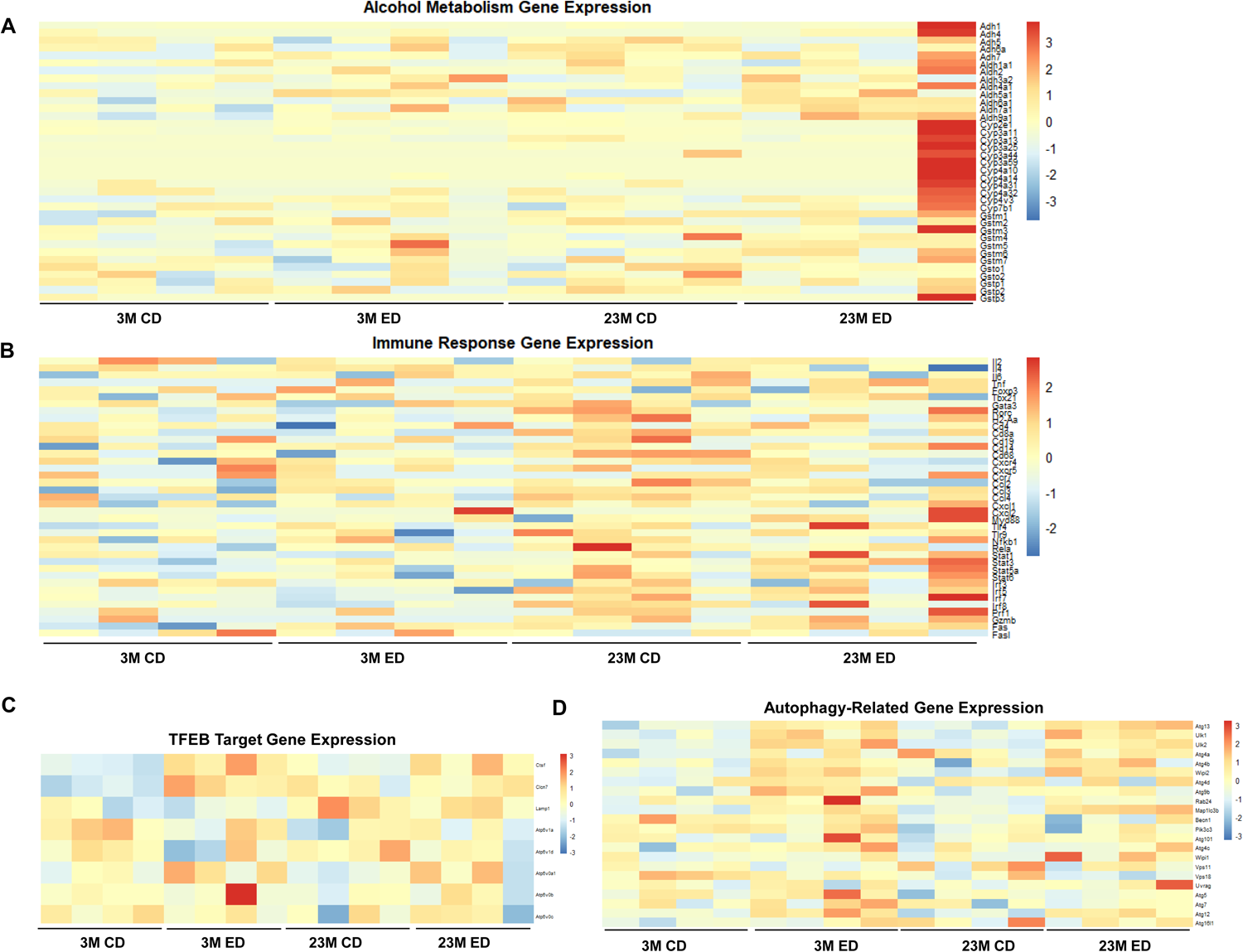
Heatmap analysis from the RNAseq dataset of Gao-binge alcohol fed young and aged mouse hippocampi. Male 3-month-old young mice (3M) and 23-month-old aged mice (23M) were subjected to Gao-binge alcohol feeding. mRNAs were extracted from the mouse hippocampi after the feeding and subjected to RNAseq analysis. The color scale represents the magnitude of gene expression, with warmer colors (red) indicating higher expression and cooler colors (blue) indicating lower expression. The x-axis represents different experimental condition, while the y-axis corresponds to the genes of interest in different pathway. The heatmap was generated using R with the ggplot2 package. (A) Heatmap analysis of alcohol metabolism gene expression from the RNAseq dataset. (B) Heatmap analysis of immune response gene expression from the RNAseq dataset. Heatmap analysis of TFEB target genes (C) and autophagy genes (D) from the RNAseq dataset.

### Chronic alcohol feeding increases cytosolic retention of TFEB in young and aged mouse brains but does not significantly affect the levels of phosphorylated Tau and histology changes in mouse brains

We next determined whether chronic ethanol feeding alone (EtOH) for 4 weeks would affect brain TFEB and AD-related markers in young and aged mice. IHC staining for TFEB revealed increased cytosolic TFEB in alcohol fed young and aged mouse cortex, but only increased in alcohol-fed aged mouse hippocampi concurrent with decreased nuclear staining (**Figure 5A-B**). Results from the western blot analysis showed that chronic alcohol feeding alone had little effect on levels of phosphorylated Tau and p62 in both young and aged mouse hippocampi (**Figure 6A-E**). The levels of ubiquitinated proteins were higher in aged mouse hippocampi compared with young mice fed the control diet. However, chronic alcohol feeding increased levels of ubiquitinated proteins in young but not aged mouse hippocampi (**Figure 6A & F**). Both young and aged mice consumed similar levels of ethanol during the four-week alcohol feeding (**Figure 7A**). Alcohol feeding decreased weight gain in young but not aged mice (**Figure 7B**). Moreover, H&E staining showed no obvious changes in brain histology in both young and aged mice regardless of alcohol feeding (**Figure 7C**). Together, these data suggest that chronic alcohol feeding increases cytosolic TFEB retention but does not impact brain histology or AD-related markers in young and aged mice.

**Figure 5.**
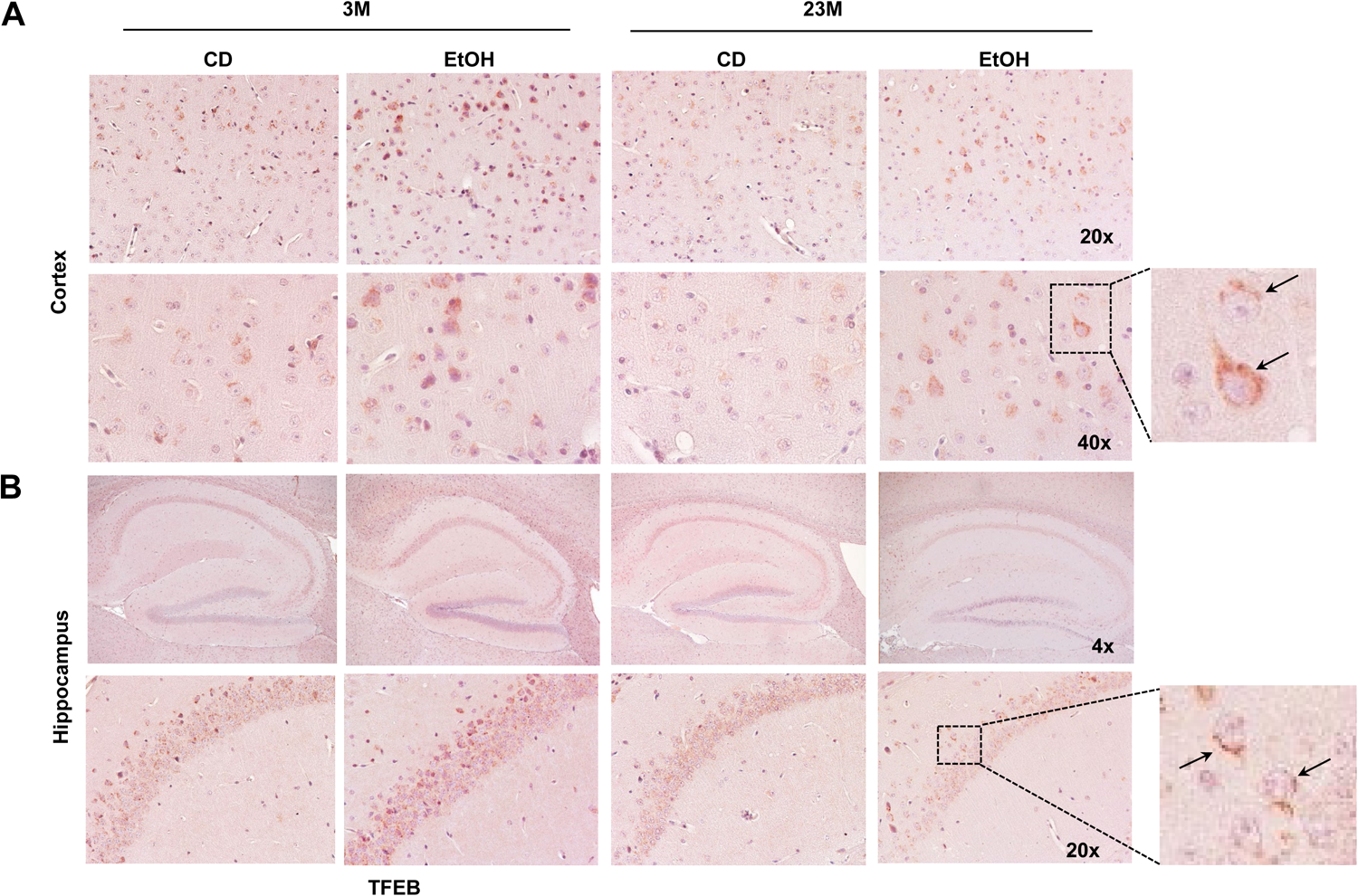
Immunohistochemistry staining of cortices and hippocampi TFEB in chronic EtOH-fed young mice and aged mice. Male 3-month-old young mice (3M) and 23-month-old aged mice (23M) were subjected to chronic EtOH feeding for four weeks. **(A)** Representative images of TFEB IHC staining of mouse cortices and (**B**) hippocampi are shown. Arrows denote cytosolic TFEB staining.

**Figure 6.**
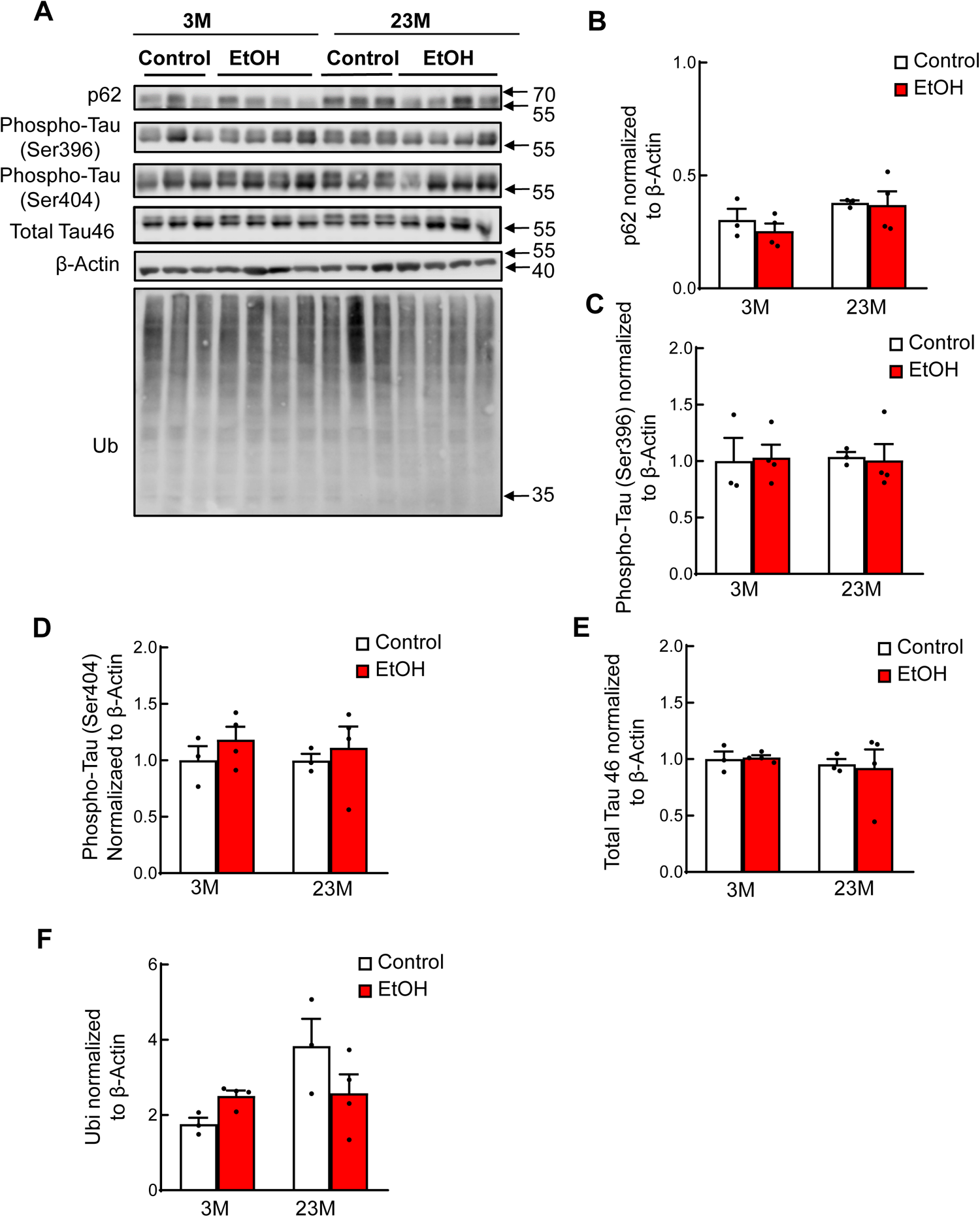
Chronic EtOH feeding does not affect Tau levels but increases levels of protein ubiquitination in young but not in aged mouse brains. Male 3-month-old young mice and 23-month-old aged mice were subjected to chronic EtOH feeding for 4 weeks. **(A)** Cortex brain lysates were subjected to western blot analysis followed by **(B-G)** Densitometry analysis of **(A),** which are normalized to loading control β-actin. Data are presented as means ± SEM (n = 3-4).

**Figure 7.**
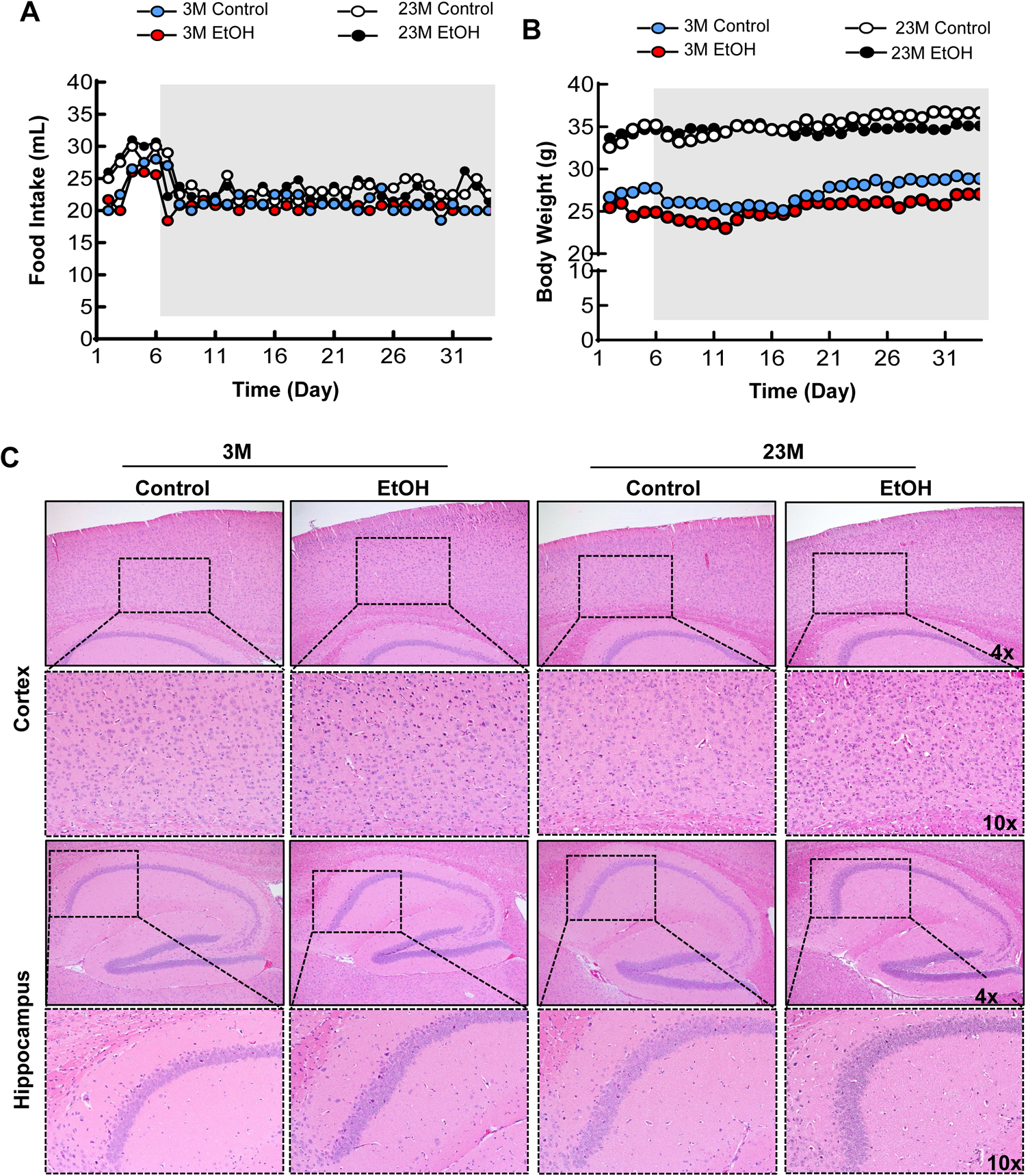
Chronic EtOH feeding increases apoptotic-like cell populations in young mice but not in aged mice brains, and without affecting body weight and food intake. Male 3-month-old young mice and 23-month-old aged mice were subjected to Gao-binge alcohol feeding. **(A)** Food intake and body weight (BW) **(B)** food intake was measured (n=4-6). **(C)** Representative brain hematoxylin and eosin images are shown. Lower panels are enlarged images from the boxed areas. Original magnifications (4x and 10x).

### Aging Increases Synaptic Loss

Synaptic loss and neurofibrillary pathology in the limbic system and neocortex contribute to AD cognitive decline (Mirza and Zahid 2018). A marked loss of the presynaptic markers synapsin-1 and synaptophysin, as well as the postsynaptic marker PSD-95, is observed in individuals with mild cognitive impairment and is associated with both prodromal and advanced stages of AD. (Scheff and Price 2001, Scheff, Price et al. 2015). Chronic alcohol feeding significantly reduced levels of hippocampal PSD95 and Synapsin. Additionally, PSD95 and Synapsin were lower in aged mice (33-months old) than they were in young mice (3-months old). Interestingly, chronic alcohol consumption did not affect the levels of these proteins in the hippocampi of the aged mice (**Figure 8A-B**).

**Figure 8.**
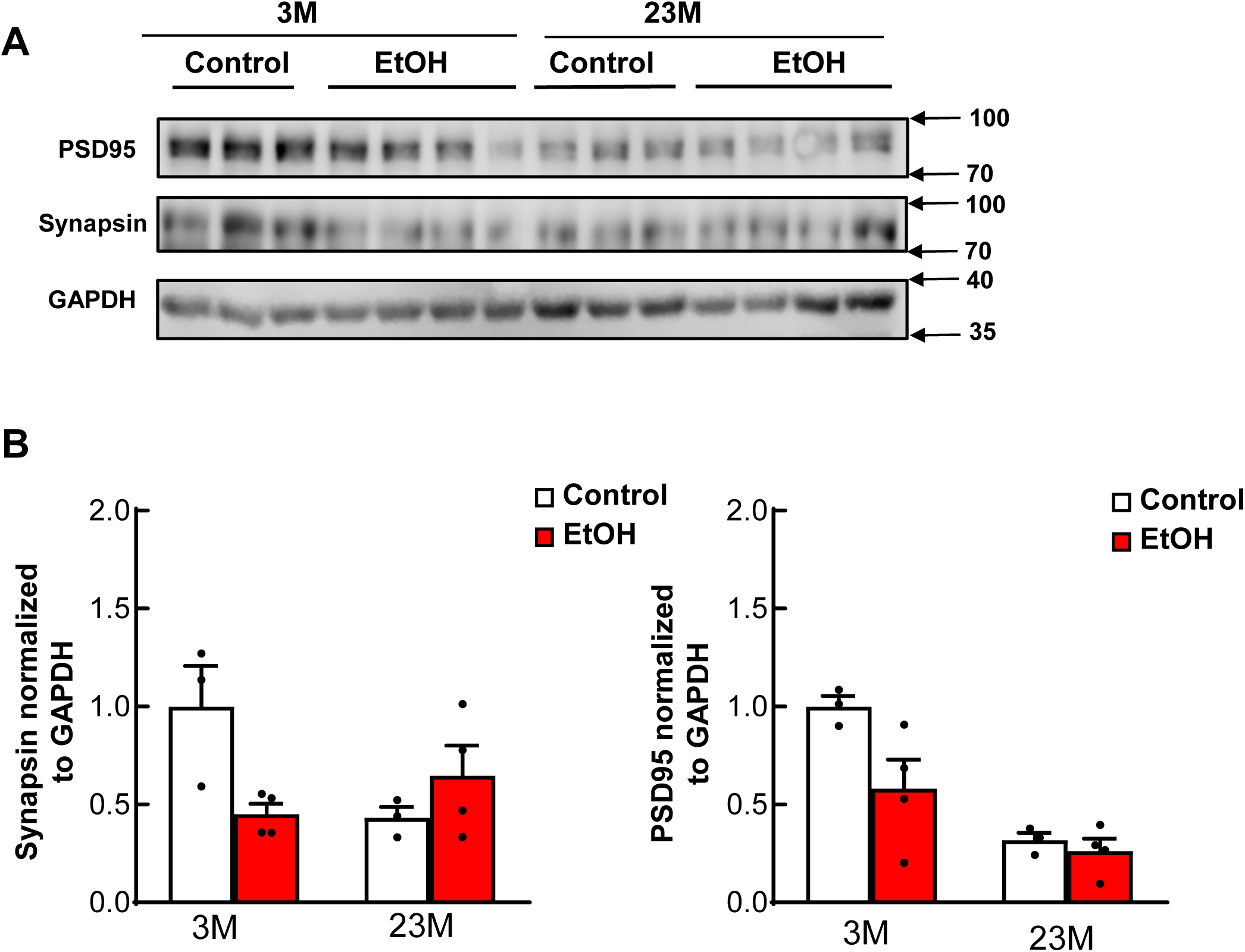
Chronic ethanol feeding has no effects on synapses but synapses are reduced in aged mice compared to young mice. Male 3-month-old young mice (3M) and 23-month-old aged mice (23M) were subjected to chronic EtOH feeding for four weeks. **(A)** Cortex brain lysates were subjected to western blot analysis followed by **(B)** Densitometry analysis of **(A),** which are normalized to loading control GAPDH. Data are presented as means ± SEM (n = 3-4) in **(B).** *p*-value; one-way analysis of variance analysis with Bonferroni’s post hoc test.

### Aging Affects Spatial Memory to a Greater Extent than Alcohol

The Morris Water Maze test assesses reference and working memory (Morris, Garrud et al. 1982, Vorhees and Williams 2006). Spatial learning was demonstrated by a decreased latency in finding the escape platform and an increased efficiency in reaching the platform. Neither aging nor alcohol feeding affected the swimming speed (**Figure 9A**), suggesting motor neurons and locomotive functions of the mice were not compromised by either advanced age or alcohol feeding. Swimming path analysis and quantification of the latency to platform time showed that aged mice fed the control diet had increased latency to finding the platform compared with young mice fed the control diet (**Figure 9B-C**). Aged mice fed the control diet also showed a trend of reduced time spent in the target quadrant (**Figure 9D**), and significantly fewer target annulus crossovers (**Figure 9E**) compared with young mice fed the control diet. Interestingly, alcohol fed young but not aged mice had increased latency to platform and decreased time spent in target and numbers of target annulus crossovers compared to their respective control diet fed mice (**Figure 9C-E**).

**Figure 9.**
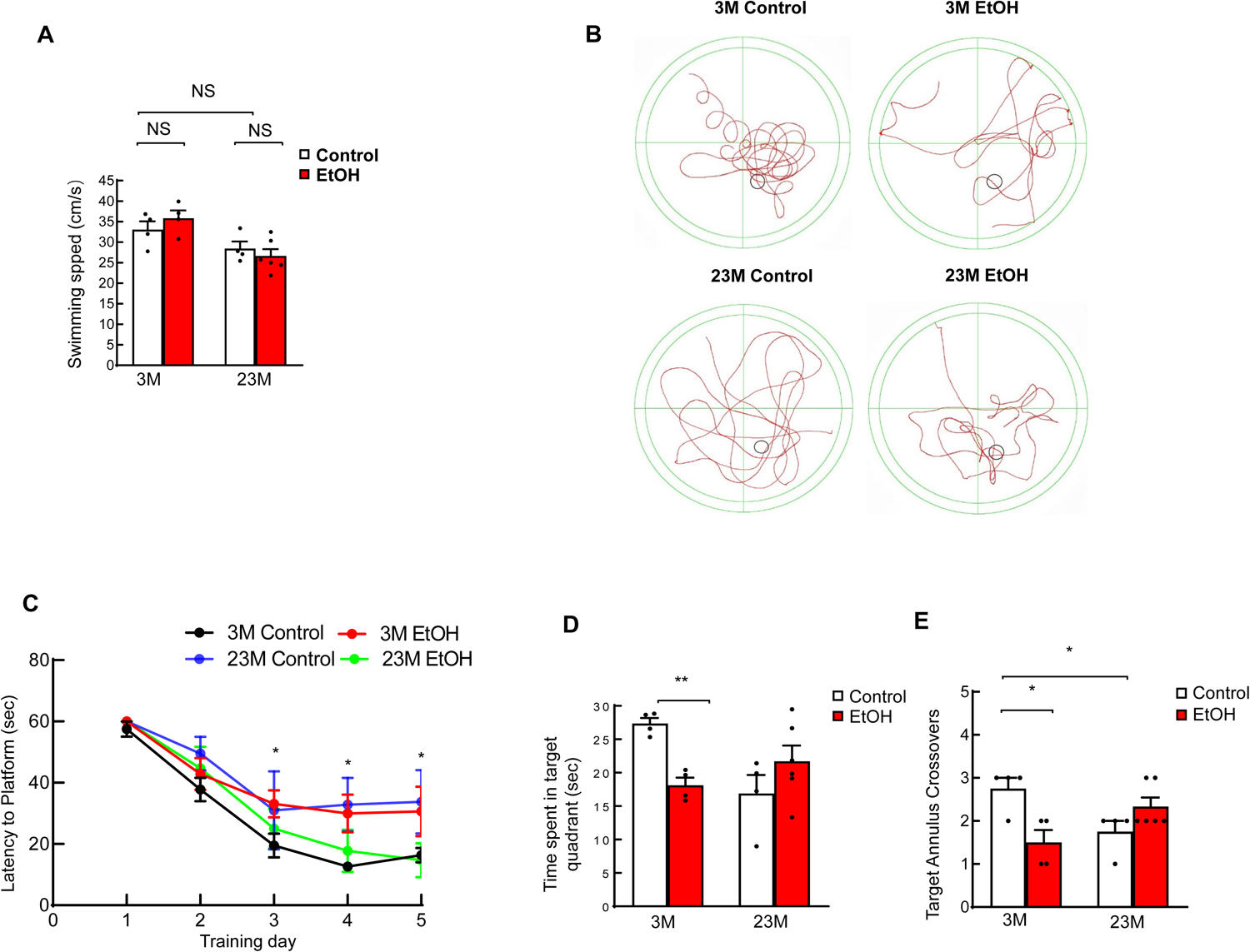
Chronic EtOH feeding impairs spatial learning and memory of young mice but not aged mice in Morris Water Maze test. Male 3-month-old young mice (3M) and 23-month-old aged mice (23M) were subjected to chronic EtOH feeding for four weeks, followed by Morris water maze test. **(A)** Swimming speed of indicated four mouse groups. **(B)** Representative images of swimming paths of indicated four mice groups during the probe test. **(C)** Escape latency decreased across the training days, representing spatial learning ability of four groups of young and aged mice fed with or without EtOH. 3M control vs 23M control. *\*p<0.05;* one-way analysis of variance analysis with Bonferroni’s post hoc test. **(D)** Spatial reference memory of indicated four mice groups evaluated by time spent in target quadrant and **(E)** number of times crossing the target platform. Data are presented as mean ± SEM. 3-month-old mice: Control, *n =* 4; EtOH, *n =* 4; 23-month-old mice: Control, *n =* 4; EtOH, *n =* 6; *\*p<0.05; **p<0.01*; one-way analysis of variance analysis with Bonferroni’s post hoc test.

The Barnes Maze test is a dry-land based behavioral test that is used to study rodent spatial memory (Barnes 1979). From a behavior interrogation perspective, it is similar to the Morris Water Maze. Heat map analysis and quantification of the latency to escape revealed that aged mice fed the control diet had increased latency of time to escape compared with young mice fed the control diet (**Figure 10A-B**), which is consistent with the Morris water maze test results, supporting the conclusion that aging impairs mouse spatial memory. Aged mice also showed an increased average distance to goal compared with the young mice, which was not affected by alcohol feeding (**Figure 10A-G**). Alcohol-fed young mice had no significant effects on latency to goal and distance to escape when compared to the control diet fed young mice (**Figure 10D-G**). Together, these data indicate that aging robustly drives reductions in spatial memory performance.

**Figure 10.**
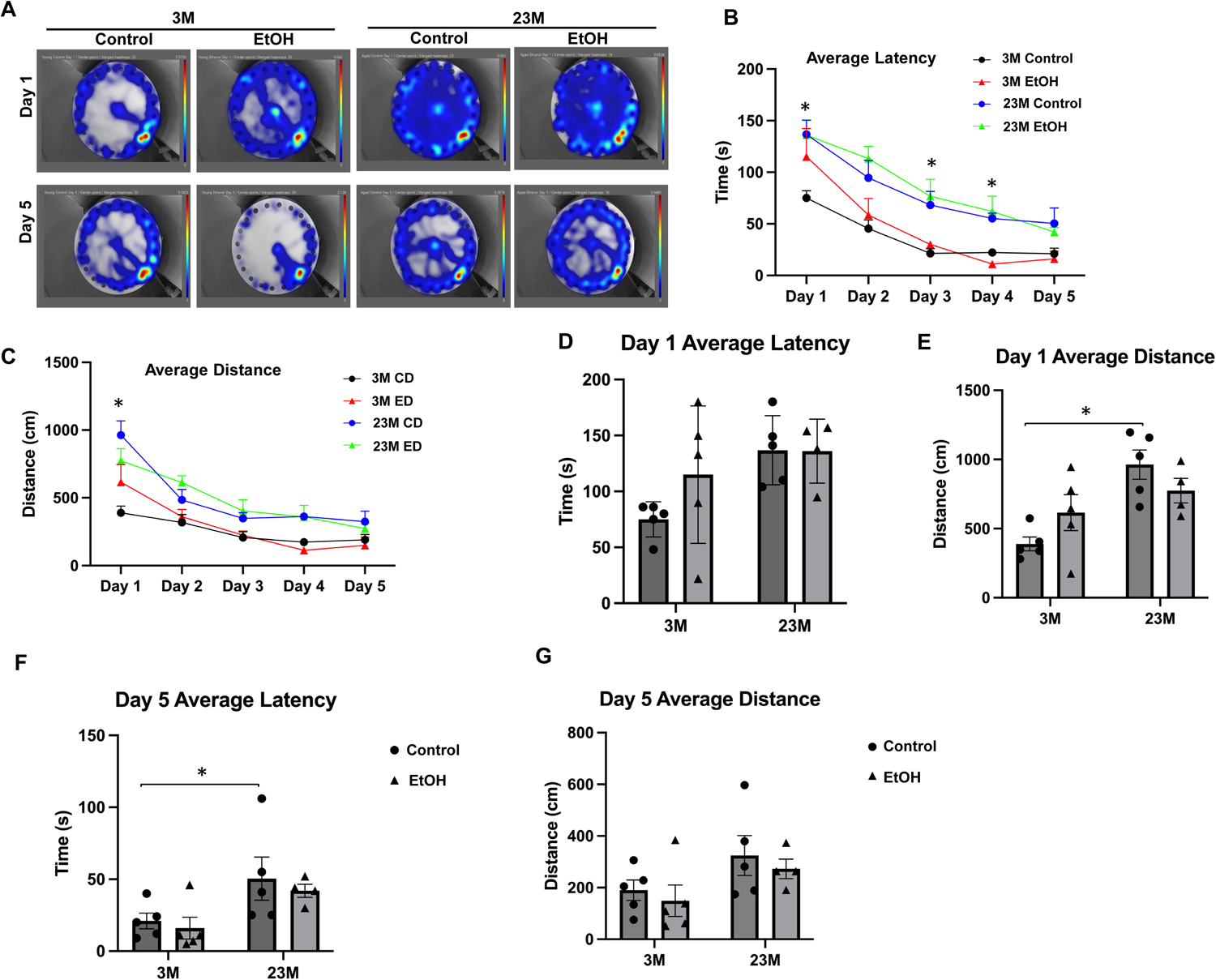
Aging but not chronic EtOH feeding impairs mouse spatial learning and memory in Barnes Maze test. Male 3-month-old young mice (3M) and 23-month-old aged mice (23M) were subjected to chronic EtOH feeding for four weeks, followed by Barnes Maze testing. **(A)** Representative heat maps of Days 1 and 5 from each treatment group. **(B)** The **a**verage latency to escape for each test day. **(C)** Average distance traveled to escape for young and aged mice fed with and without EtOH. **(D)** Day 1 average latency to escape for 3-month- and 23-month-old mice fed with or without EtOH. 3M control vs 23M control (B-C). *\*p<0.05;* one-way analysis of variance analysis with Dunn’s post hoc test. **(E)** Day 1 average distance traveled to escape for 3-month- and 23-month-old mice fed with or without EtOH. **(F)** Day 5 average latency to escape for 3-month- and 23-month-old mice fed with or without EtOH. **(G)** Day 5 average distance traveled to escape for 3-month- and 23-month-old mice fed with or without EtOH. Data are presented as mean ± SEM. 3-month-old: CD, n = 5; ED, n = 5; 23-month-old: CD, n = 5; ED, n = 4. *\*p<0.05;* one-way analysis of variance analysis with Dunn’s post hoc test.

## Discussion

In this study, we used two alcohol-feeding models to address the impact of alcohol and aging on brain autophagy and TFEB as well as mouse cognitive activities. We used the Gao-binge alcohol model, which is a widely used to mimics chronic drinking plus binge alcohol consumption (Bertola, Mathews et al. 2013). Biochemical, transcriptomic, and histologic analysis of hippocampi from young and aged mice showed impaired TFEB-mediated autophagy in Gao-binge alcohol-fed young mice. Brains from aged mice showed defective TFEB-mediated autophagy relative to brains from young mice, and Gao-binge alcohol feeding did not exacerbate the age-related changes. Aged mice had impaired spatial memory in the Morris Water Maze and Barnes Maze tests. While alcohol feeding slightly impaired spatial memory in young mice, alcohol had little effect or even slightly improved spatial memory in aged mice.

AD is a neurodegenerative condition that is primarily associated with the abnormal processing and polymerization of normally soluble proteins. Gene mutations and aging can promote protein misfolding and aggregation, in which associates with neuron dysfunction and loss. AD is generally characterized by extracellular aggregates of Aβ plaques and intracellular aggregations of neurofibrillary tangles (NFTs) composed of hyperphosphorylated tau protein. In addition to Aβ plaques and tau aggregation, mitochondrial dysfunction also associates with the disease (Ashleigh, Swerdlow et al. 2023). Increasing evidence suggests impaired autophagy may cause an accumulation of Aβ and tau aggregates and contribute to the development of AD (Zhang, Chen et al. 2023). Numerous studies show autophagy is disturbed in AD patients, and potentially drives an accumulation of autophagic vehicles (AVs) within neurons (Terry, Gonatas et al. 1964, Nixon, Wegiel et al. 2005). Lysosomal biogenesis is regulated by TFEB. Dysfunctional TFEB has been linked to AD in both humans and experimental AD models (Wang, Wang et al. 2016, Zhang, Chen et al. 2023). Moreover, in AD transgenic mouse models, pharmacological activation or genetic overexpression of TFEB initiates a lysosomal biogenesis that clears Aβ and tau aggregates (Polito, Li et al. 2014, Wang, Wang et al. 2016, Song, Malampati et al. 2020). Aging is a well-known AD risk factor, and aging associates with reductions in autophagy and TFEB (Rubinsztein, Mariño et al. 2011, Wang, Karuppan et al. 2021).

We found aging alone slightly decreased brain TFEB levels with increased LC3-II, p62 and ubiquitination by western blot and IHC analysis, and alcohol feeding did not enhance these changes. Moreover, our transcriptome analysis also showed decreased expression of some TFEB target genes and autophagy genes in aged mouse brains, and alcohol exposure did not exacerbate these changes. This suggests aging more robustly reduces brain TFEB-mediated autophagy than alcohol.

Morris Water and Barnes Maze testing revealed spatial memory in aged mice was impaired relative to young mice. Alcohol impaired spatial memory in Morris Water Maze but not Barnes Maze testing in young mice. Surprisingly, alcohol feeding did not worsen the age-related spatial memory decrements in aged mice. Our biochemical, transcriptome and histological observations are generally in line with the results from the two spatial memory tests. Aged mice already have lower levels of TFEB in their brains, regardless of whether they consumed alcohol or not, while alcohol decreased TFEB in younger mice. Our findings support the notion that age-related reductions in autophagy and TFEB could contribute to the development of AD. C1, a curcumin analog, efficiently activated TFEB, enhanced autophagy and lysosomal activity, and reduced β-amyloid peptides and Tau aggregates in these models, which was accompanied by improved synaptic and cognitive function (Song, Malampati et al. 2020). However, we failed to observe increased TFEB expression in brains from mice treated with C1 and alcohol (data not shown). Further studies are needed to address whether TFEB is a valid therapeutic target (Chao, Niu et al. 2024).

Several possibilities could account for why alcohol did not worsen cognitive performance in aged mice. The duration of exposure was only four weeks. A more chronic treatment may have produced a different result. During the treatment period we observed the alcohol-fed mice increased their brain expression of alcohol metabolism-related genes. This suggests they mounted an acute adaptive response. We don’t know whether this adaptive response persists under conditions of chronic alcohol exposure.

The impact of alcohol consumption on AD risk remains unclear. Inconsistent results from past studies may reflect inter-study differences in how consumption amounts and patterns were defined (Tyas 2001, Heymann, Stern et al. 2016). Some studies indicate moderate alcohol consumption may reduce AD risk in older populations (Neafsey and Collins 2011, Weyerer, Schäufele et al. 2011). Additional research is needed to clarify the impact of alcohol on AD risk and pathogenesis.

## Author contributions

H.C., K.H., C.Z., Y.R., S.W., M. N., X.M., X.C., A.L.F., K.M. and H.M.N. performed the experiments and analyzed data. C.Z., P.Z. and W.L performed RNAseq data analysis and interpretation. W.X.D., H.C. & H.M.N conceived the idea and wrote the manuscript. K.M., J.Z. & R.H.S. assisted experiment design and revised the manuscript.

## Abbreviations

AD: Alzheimer’s disease
ADRD: AD and its related dementia
AVs: autophagic vehicles
CLEAR: coordinated lysosomal expression and regulation
CD: Control diet
CNS: *c*entral nervous system
ED: ethanol diet
*Fkbp5*: FK506-binding protein 5
HPA: hypothalamic-pituitary-adrenal
LD: lipid droplets
NFTs: neurofibrillary tangles
PTSD: posttraumatic stress disorder
SNP: single nucleotide polymorphism
TFEB: Transcription factor EB

## References

Androuin, A., M. Thierry, S. Boluda, A. Baskaran, D. Langui, C. Duyckaerts, M. C. Potier, K. H. El Hachimi, B. Delatour, S. Marty and B. N. Neuropathology (2022). “Alterations of Neuronal Lysosomes in Alzheimer’s Disease and in APPxPS1-KI Mice.” Journal of Alzheimers Disease 87(1): 273–284.

Ashleigh, T., R. H. Swerdlow and M. F. Beal (2023). “The role of mitochondrial dysfunction in Alzheimer’s disease pathogenesis.” Alzheimers & Dementia 19(1): 333–342.

Babuta, M., I. Furi, S. Bala, T. N. Bukong, P. Lowe, D. Catalano, C. Calenda, K. Kodys and G. Szabo (2019). “Dysregulated Autophagy and Lysosome Function Are Linked to Exosome Production by Micro-RNA 155 in Alcoholic Liver Disease.” Hepatology 70(6): 2123–2141.

Bao, J., L. Zheng, Q. Zhang, X. Li, X. Zhang, Z. Li, X. Bai, Z. Zhang, W. Huo, X. Zhao, S. Shang, Q. Wang, C. Zhang and J. Ji (2016). “Deacetylation of TFEB promotes fibrillar Abeta degradation by upregulating lysosomal biogenesis in microglia.” Protein Cell 7(6): 417–433.

Barnes, C. A. (1979). “Memory deficits associated with senescence: a neurophysiological and behavioral study in the rat.” J Comp Physiol Psychol 93(1): 74–104.

Bertola, A., S. Mathews, S. H. Ki, H. Wang and B. Gao (2013). “Mouse model of chronic and binge ethanol feeding (the NIAAA model).” Nat Protoc 8(3): 627–637.

Chan, K. K. K., K. C. Chiu and L. W. Chu (2010). “Association between alcohol consumption and cognitive impairment in Southern Chinese older adults.” International Journal of Geriatric Psychiatry 25(12): 1272–1279.

Chao, X., M. Niu, S. Wang, X. Ma, X. Yang, H. Sun, X. Hu, H. Wang, L. Zhang, R. Huang, M. Xia, A. Ballabio, H. Jaeschke, H. M. Ni and W. X. Ding (2024). “High-throughput screening of novel TFEB agonists in protecting against acetaminophen-induced liver injury in mice.” Acta Pharm Sin B 14(1): 190–206.

Chao, X., S. Wang, K. Zhao, Y. Li, J. A. Williams, T. Li, H. Chavan, P. Krishnamurthy, X. C. He, L. Li, A. Ballabio, H. M. Ni and W. X. Ding (2018). “Impaired TFEB-Mediated Lysosome Biogenesis and Autophagy Promote Chronic Ethanol-Induced Liver Injury and Steatosis in Mice.” Gastroenterology 155(3): 865–879 e812.

Czaja, M. J., W. X. Ding, T. M. Donohue, Jr., S. L. Friedman, J. S. Kim, M. Komatsu, J. J. Lemasters, A. Lemoine, J. D. Lin, J. H. Ou, D. H. Perlmutter, G. Randall, R. B. Ray, A. Tsung and X. M. Yin (2013). “Functions of autophagy in normal and diseased liver.” Autophagy 9(8): 1131–1158.

Fujii, T., H. Hori, M. Ota, K. Hattori, T. Teraishi, D. Sasayama, N. Yamamoto, T. Higuchi and H. Kunugi (2014). “Effect of the common functional variant (rs1360780) on the hypothalamic-pituitary-adrenal axis and peripheral blood gene expression.” Psychoneuroendocrinology 42: 89–97.

Fujii, T., M. Ota, H. Hori, K. Hattori, T. Teraishi, J. Matsuo, Y. Kinoshita, I. Ishida, A. Nagashima and H. Kunugi (2014). “The common functional variant rs1360780 is associated with altered cognitive function in aged individuals.” Scientific Reports 4.

Hardy, J. (2006). “A hundred years of Alzheimer’s disease research.” Neuron 52(1): 3–13.

Heymann, D., Y. Stern, S. Cosentino, O. Tatarina-Nulman, J. N. Dorrejo and Y. Gu (2016). “The Association Between Alcohol Use and the Progression of Alzheimer’s Disease.” Current Alzheimer Research 13(12): 1356–1362.

Jeon, K. H., K. Han, S. M. Jeong, J. Park, J. E. Yoo, J. Yoo, J. Lee, S. Kim and D. W. Shin (2023). “Changes in Alcohol Consumption and Risk of Dementia in a Nationwide Cohort in South Korea.” Jama Network Open 6(2).

Klionsky, D. J. and S. D. Emr (2000). “Autophagy as a regulated pathway of cellular degradation.” Science 290(5497): 1717–1721.

Levine, B. and G. Kroemer (2019). “Biological Functions of Autophagy Genes: A Disease Perspective.” Cell 176(1-2): 11–42.

Ma, X., A. Chen, L. Melo, A. Clemente-Sanchez, X. Chao, A. R. Ahmadi, B. Peiffer, Z. Sun, H. Sesaki, T. Li, X. Wang, W. Liu, R. Bataller, H. M. Ni and W. X. Ding (2023). “Loss of hepatic DRP1 exacerbates alcoholic hepatitis by inducing megamitochondria and mitochondrial maladaptation.” Hepatology 77(1): 159–175.

Martini-Stoica, H., Y. Xu, A. Ballabio and H. Zheng (2016). “The Autophagy-Lysosomal Pathway in Neurodegeneration: A TFEB Perspective.” Trends Neurosci 39(4): 221–234.

Mirza, F. J. and S. Zahid (2018). “The Role of Synapsins in Neurological Disorders.” Neuroscience Bulletin 34(2): 349–358.

Mizushima, N., B. Levine, A. M. Cuervo and D. J. Klionsky (2008). “Autophagy fights disease through cellular self-digestion.” Nature 451(7182): 1069–1075.

Morris, R. G., P. Garrud, J. N. Rawlins and J. O’Keefe (1982). “Place navigation impaired in rats with hippocampal lesions.” Nature 297(5868): 681–683.

Neafsey, E. J. and M. A. Collins (2011). “Moderate alcohol consumption and cognitive risk.” Neuropsychiatric Disease and Treatment 7: 465–484.

Nixon, R. A. (2007). “Autophagy, amyloidogenesis and Alzheimer disease.” Journal of Cell Science 120(23): 4081–4091.

Nixon, R. A., J. Wegiel, A. Kumar, W. H. Yu, C. Peterhoff, A. Cataldo and A. M. Cuervo (2005). “Extensive involvement of autophagy in Alzheimer disease: an immuno-electron microscopy study.” J Neuropathol Exp Neurol 64(2): 113–122.

Pitts, M. W. (2018). “Barnes Maze Procedure for Spatial Learning and Memory in Mice.” Bio Protoc 8(5).

Piumatti, G., S. C. Moore, D. M. Berridge, C. Sarkar and J. Gallacher (2018). “The relationship between alcohol use and long-term cognitive decline in middle and late life: a longitudinal analysis using UK Biobank.” J Public Health (Oxf) 40(2): 304–311.

Polito, V. A., H. Li, H. Martini-Stoica, B. Wang, L. Yang, Y. Xu, D. B. Swartzlander, M. Palmieri, A. di Ronza, V. M. Lee, M. Sardiello, A. Ballabio and H. Zheng (2014). “Selective clearance of aberrant tau proteins and rescue of neurotoxicity by transcription factor EB.” EMBO Mol Med 6(9): 1142–1160.

Reddy, K., C. L. Cusack, I. C. Nnah, K. Khayati, C. Saqcena, T. B. Huynh, S. A. Noggle, A. Ballabio and R. Dobrowolski (2016). “Dysregulation of Nutrient Sensing and CLEARance in Presenilin Deficiency.” Cell Rep 14(9): 2166–2179.

Rodriguez Peris, L., M. I. Scheuber, H. Shan, M. Braun and M. E. Schwab (2024). “Barnes maze test for spatial memory: A new, sensitive scoring system for mouse search strategies.” Behav Brain Res 458: 114730.

Rubinsztein, D. C., G. Mariño and G. Kroemer (2011). “Autophagy and Aging.” Cell 146(5): 682–695.

Scheff, S. W. and D. A. Price (2001). “Alzheimer’s disease-related synapse loss in the cingulate cortex.” Journal of Alzheimers Disease 3(5): 495–505.

Scheff, S. W., D. A. Price, M. A. Ansari, K. N. Roberts, F. A. Schmitt, M. D. Ikonomovic and E. J. Mufson (2015). “Synaptic Change in the Posterior Cingulate Gyrus in the Progression of Alzheimer’s Disease.” Journal of Alzheimers Disease 43(3): 1073–1090.

Schwarzinger, M., B. G. Pollock, O. S. M. Hasan, C. Dufouil, J. Rehm and G. QalyDays Study (2018). “Contribution of alcohol use disorders to the burden of dementia in France 2008-13: a nationwide retrospective cohort study.” Lancet Public Health 3(3): e124–e132.

Settembre, C., C. Di Malta, V. A. Polito, M. Garcia Arencibia, F. Vetrini, S. Erdin, S. U. Erdin, T. Huynh, D. Medina, P. Colella, M. Sardiello, D. C. Rubinsztein and A. Ballabio (2011). “TFEB links autophagy to lysosomal biogenesis.” Science 332(6036): 1429–1433.

Settembre, C., A. Fraldi, D. L. Medina and A. Ballabio (2013). “Signals from the lysosome: a control centre for cellular clearance and energy metabolism.” Nat Rev Mol Cell Biol 14(5): 283–296.

Settembre, C., R. Zoncu, D. L. Medina, F. Vetrini, S. Erdin, S. Erdin, T. Huynh, M. Ferron, G. Karsenty, M. C. Vellard, V. Facchinetti, D. M. Sabatini and A. Ballabio (2012). “A lysosome-to-nucleus signalling mechanism senses and regulates the lysosome via mTOR and TFEB.” EMBO J 31(5): 1095–1108.

Singh, R., S. Kaushik, Y. Wang, Y. Xiang, I. Novak, M. Komatsu, K. Tanaka, A. M. Cuervo and M. J. Czaja (2009). “Autophagy regulates lipid metabolism.” Nature 458(7242): 1131–1135.

Song, J. X., S. Malampati, Y. Zeng, S. S. K. Durairajan, C. B. Yang, B. C. Tong, A. Iyaswamy, W. B. Shang, S. G. Sreenivasmurthy, Z. Zhu, K. H. Cheung, J. H. Lu, C. Tang, N. Xu and M. Li (2020). “A small molecule transcription factor EB activator ameliorates beta-amyloid precursor protein and Tau pathology in Alzheimer’s disease models.” Aging Cell 19(2): e13069.

Song, J. X., S. Malampati, Y. Zeng, S. S. K. Durairajan, C. B. Yang, B. C. K. Tong, A. Iyaswamy, W. B. Shang, S. G. Sreenivasmurthy, Z. Zhu, K. H. Cheung, J. H. Lu, C. Z. Tang, N. G. Xu and M. Li (2020). “A small molecule transcription factor EB activator ameliorates beta-amyloid precursor protein and Tau pathology in Alzheimer’s disease models.” Aging Cell 19(2).

Stavro, K., J. Pelletier and S. Potvin (2013). “Widespread and sustained cognitive deficits in alcoholism: a meta-analysis.” Addict Biol 18(2): 203–213.

Swerdlow, R. H. (2018). “Mitochondria and Mitochondrial Cascades in Alzheimer’s Disease.” J Alzheimers Dis 62(3): 1403–1416.

Terry, R. D., N. K. Gonatas and M. Weiss (1964). “The ultrastructure of the cerebral cortex in Alzheimer’s disease.” Trans Am Neurol Assoc 89: 12.

Tiribuzi, R., L. Crispoltoni, S. Porcellati, M. Di Lullo, F. Florenzano, M. Pirro, F. Bagaglia, T. Kawarai, M. Zampolini, A. Orlacchio and A. Orlacchio (2014). “miR128 up-regulation correlates with impaired amyloid beta(1-42) degradation in monocytes from patients with sporadic Alzheimer’s disease.” Neurobiol Aging 35(2): 345–356.

Tyas, S. L. (2001). “Alcohol use and the risk of developing Alzheimer’s disease.” Alcohol Research & Health 25(4): 299–306.

Van Giau, V., S. S. A. An and J. P. Hulme (2018). “Mitochondrial therapeutic interventions in Alzheimer’s disease.” J Neurol Sci 395: 62–70.

Vorhees, C. V. and M. T. Williams (2006). “Morris water maze: procedures for assessing spatial and related forms of learning and memory.” Nat Protoc 1(2): 848–858.

Wang, H., R. Wang, I. Carrera, S. Xu and M. K. Lakshmana (2016). “TFEB Overexpression in the P301S Model of Tauopathy Mitigates Increased PHF1 Levels and Lipofuscin Puncta and Rescues Memory Deficits.” eNeuro 3(2).

Wang, H., R. Wang, S. Xu and M. K. Lakshmana (2016). “Transcription Factor EB Is Selectively Reduced in the Nuclear Fractions of Alzheimer’s and Amyotrophic Lateral Sclerosis Brains.” Neurosci J 2016: 4732837.

Wang, H. J., M. K. M. Karuppan, D. Devadoss, M. Nair, H. S. Chand and M. K. Lakshmana (2021). “TFEB protein expression is reduced in aged brains and its overexpression mitigates senescence-associated biomarkers and memory deficits in mice.” Neurobiology of Aging 106: 26–36.

Wang, S., H. M. Ni, X. Chao, X. Ma, T. Kolodecik, R. De Lisle, A. Ballabio, P. Pacher and W. X. Ding (2020). “Critical Role of TFEB-Mediated Lysosomal Biogenesis in Alcohol-Induced Pancreatitis in Mice and Humans.” Cell Mol Gastroenterol Hepatol.

Weyerer, S., M. Schäufele, B. Wiese, W. Maier, F. Tebarth, H. van den Bussche, M. Pentzek, H. Bickel, M. Luppa, S. G. Riedel-Heller and G. A. S. Grp (2011). “Current alcohol consumption and its relationship to incident dementia: results from a 3-year follow-up study among primary care attenders aged 75 years and older.” Age and Ageing 40(4): 456–463.

Williams, J. A., H. M. Ni, Y. Ding and W. X. Ding (2015). “Parkin regulates mitophagy and mitochondrial function to protect against alcohol-induced liver injury and steatosis in mice.” Am J Physiol Gastrointest Liver Physiol 309(5): G324–340.

Zhang, C., H. Chen, Y. Rodriguez, X. W. Ma, R. H. Swerdlow, J. H. Zhang and W. X. Ding (2023). “A perspective on autophagy and transcription factor EB in Alcohol-Associated Alzheimer’s disease.” Biochemical Pharmacology 213.

